# Performance of temporal and spatial ICA in identifying and removing low-frequency physiological and motion effects in resting-state fMRI

**DOI:** 10.1101/2021.09.19.460965

**Authors:** Ali M Golestani, J. Jean Chen

## Abstract

Effective separation of signal from noise (including physiological processes and head motion) is one of the chief challenges for improving the sensitivity and specificity of resting-state fMRI (rs-fMRI) measurements and has a profound impact when these noise sources vary between populations. Independent component analysis (ICA) is an approach for addressing these challenges. Conventionally, due to the lower amount of temporal than spatial information in rs-fMRI data, spatial ICA (sICA) is the method of choice. However, with recent developments in accelerated fMRI acquisitions, the temporal information is becoming enriched to the point that the temporal ICA (tICA) has become more feasible. This is particularly relevant as physiological processes and motion exhibit very different spatial and temporal characteristics when it comes to rs-fMRI applications, leading us to conduct a comparison of the performance of sICA and tICA in addressing these types of noise. In this study, we embrace the novel practice of using theory (simulations) to guide our interpretation of empirical data. We find empirically that sICA can identify more noise-related signal components than tICA. However, on the merit of functional-connectivity results, we find that while sICA is more adept at reducing whole-brain motion effects, tICA performs better in dealing with physiological effects. These interpretations are corroborated by our simulation results. The overall message of this study is that if ICA denoising is to be used for rs-fMRI, there is merit in considering a hybrid approach in which physiological and motion-related noise are each corrected for using their respective best-suited ICA approach.

**Impact Statement:** Resting-state fMRI is influenced by low-frequency physiological noise and head motion. Independent component analysis (ICA) is becoming increasingly relied on for reducing these influences, but the utility of spatial and temporal ICA remains unclear. We conducted a comparison of the performance of these two ICA types, using physiological-noise and motion time courses as reference. We found that spatial ICA is more adept at reducing motion effects, while temporal ICA performs better in dealing with physiological effects. We believe these findings provide much-needed clarity on the role of ICA, and recommend using a hybrid of tICA and sICA as a paradigm shift in resting-state fMRI.

## Introduction

Functional MRI (fMRI) is a powerful tool to non-invasively investigate brain function and organization. However, several confounding noise sources typically affect the sensitivity and specificity of associated results, chiefly coming from physiological processes and bulk head motion. These nuisance effects typically need to be removed to reduce false positive and false negative results, particularly in resting-state fMRI (rs-fMRI). Desirable clean-up methods should selectively remove noise while preserving signals of interest generated by presumed neural activity (Caballero-Gaudes & Reynolds, 2017; Murphy et al., 2013). Identifying and removing physiological noise such as those induced by temporal variability in respiratory volume and heart rate (Birn et al., 2006; Chang et al., 2009; Golestani et al., 2015), as well as by head motion (Maknojia et al., 2019; Power et al., 2012, 2015; Yan et al., 2013) are particularly challenging. While the higher-frequency respiration and cardiac cycles have been better characterized and found easier to correct (G. H. Glover et al., 2000), the low-frequency physiological effects have characteristics that vary among different subjects and populations. Moreover, subtle head motion, which is likely connected to such physiological effects in no small part, is notoriously hard to identify and remove (Van Dijk et al., 2012).

Several studies have used independent component analysis (ICA) to remove the effects of physiological signals from fMRI (Pruim et al., 2015; Salimi-Khorshidi et al., 2014). ICA is a data-driven method that can be used to identify physiological components of the fMRI signal without *a priori* knowledge about their dynamics or additional equipment to record the physiological signals. Since fMRI data typically has more voxels than time-points, so far it has been more feasible to perform spatial ICA (sICA) on fMRI data (Smith et al., 2012). Indeed, most of the available ICA-based data-cleaning tools are based on sICA (Griffanti et al., 2014; Pruim et al., 2015; Salimi-Khorshidi et al., 2014). However, recent developments in multiband data acquisition enable acquiring fMRI with higher temporal resolutions, which makes temporal ICA (tICA) more feasible (Alkan et al., 2011; Amemiya et al., 2019; Baker et al., 2019; Boubela et al., 2013; Calhoun et al., 2001; Chen et al., 2006; Gao et al., 2011; Hald et al., 2017; Lukic et al., 2007; R. L. Miller et al., 2014; Penney & Koles, 2006; Shi & Zeng, 2018; Stone et al., 2002; van de Ven et al., 2009; Z. Wang et al., 2006). tICA has begun to be used in noise identification in rs-fMRI (Beall & Lowe, 2007; Glasser et al., 2018, 2019; Power, 2019). Regardless, sICA is still the method of choice for rs-fMRI noise removal.

Previous studies indicate inherent differences in denoising performance by tICA and sICA. Although sICA can successfully identify spatially localized fluctuations, it likely fails to separate spatially global components (Glasser et al., 2018) with spatially overlapping sources (Boubela et al., 2013; Smith et al., 2012). Specifically, Calhoun et al., (Calhoun et al., 2001) have shown that sICA and tICA fail in separating underlying components if the components are spatially and temporally inter-dependent, respectively. The deficiency of sICA in fMRI clean-up has been demonstrated (Burgess et al., 2016; Siegel et al., 2017), and tICA has shown promising results in identifying physiological noise in the fMRI data. Boubela et al. (Boubela et al., 2013) were able to identify physiological signals such as cardiac pulsation using tICA. Moreover, Glasser et al. (Glasser et al., 2018) used tICA as a replacement for the controversial global signal regression and showed tICA can identify and remove global fluctuations in the fMRI data (which presumably is due to physiological nuisance) while preserving neural signals. However, in both studies, the data from multiple subjects were concatenated and a group-wise tICA on the concatenated data was performed. Therefore, it is not clear how tICA performs in terms of a single-subject ICA. Moreover, the study by Glasser et al. (Glasser et al., 2018) mainly focused on identifying a component associated with the global signal and did not investigate how tICA performed in identifying and removing specific physiological noise effects.

Intuitively, tICA can better distinguish between temporally independent but spatially overlapping components (Boubela et al., 2013; Calhoun et al., 2001; Glasser et al., 2018) compared to sICA. These include effects of low-frequency physiological fluctuations, which could encompass the well-known effects of respiratory variability (RVT), heart-rate variability (HRV), and end-tidal carbon dioxide (PETCO_2_), potentially overlapping with each other spatially and temporally (Yunjie Tong et al., 2019)(Bright et al., 2020). Some of these physiological signals are also temporally related to one another (Chang & Glover, 2009; Glasser et al., 2018; Power, Lynch, Silver, et al., 2019). On the other hand, some noise sources are spatially less restricted and overlap with the spatial pattern of other noises as well as with several resting-state networks (Golestani et al., 2015)(Griffanti et al., 2014; Salimi-Khorshidi et al., 2014) It is unclear if ICA (either temporal or spatial) can identify and separate all of these sources of noise.

In this study, we compare the performance of sICA and tICA in identifying and removing physiological noises on a single-session (non-concatenated) basis. The first objective of this study is to investigate if sICA and tICA can identify different noise components in different ICs. Since different noises have spatial or temporal dependencies, we hypothesize that tICA and sICA perform differently in identifying different physiological components of the resting-state fMRI (rs-fMRI) signal. The second objective of this study is to compare tICA and sICA performance in removing noise from the fMRI data while preserving the information about brain function. It is not immediately obvious which would excel in the preservation of neuronal information.

## Theory

### ICA

ICA is a method for decomposing multivariate linearly combined signals into its components, assuming the components are statistically independent and non-Gaussian. Assuming we observe *m* signals *X = (x*_*1*_, *…,x*_*m*_)^*T*^, which are a linear mixture of *n* hidden components *S = (s*_*1*_, *…, s*_*n*_)^*T*^. The mixture can be written as a matrix multiplication as follow:

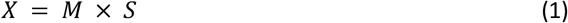

where *M* is an *m×n* mixing matrix. Assuming the components in *S* are statistically independent, ICA tries to estimate a separating matrix *W* so that

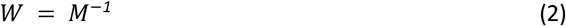

Using *W*, we can estimate the original components:

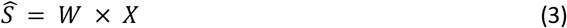

### Spatial ICA

To implement spatial ICA (Beckmann et al., 2005) on fMRI data, the data is first reordered into a 2-dimensional matrix of time x space. Assuming we have *n* voxels with *t* time samples, the fMRI data can be modeled as:

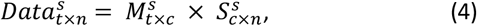

where *S*^*s*^ is a *c×n* matrix of *c* spatially independent components and *M*^*s*^ is a mixing matrix that consists of temporal signatures of the spatial components. Note that in this case we assume that the data consists of *c* spatially independent components. Using sICA we estimate the separating matrix as

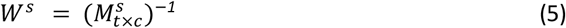

Using the separating matrix, we estimate the spatial components 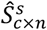:

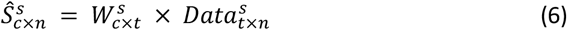

Time courses of the components can be estimated by calculating the pseudo-inverse of the separating matrix *W*^*s*^.

### Temporal ICA

To perform temporal ICA (Boubela et al., 2013; Glasser et al., 2018; Salimi-Khorshidi et al., 2014; Smith et al., 2012), the fMRI data is transposed into a *n×t* matrix. Therefore, the data can be modeled as:

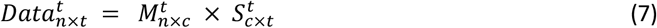

The three matrices of *Data*^*t*^, *M*^*t*^ and *S*^*t*^ are the transpose of *Data*^*s*^, *M*^*s*^ and *S*^*s*^ for the spatial ICA case. We assume that the components time series in *S*^*t*^ are independent and we try to estimate the separating matrix

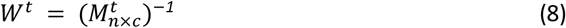

Then we can estimate the components time courses 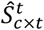:

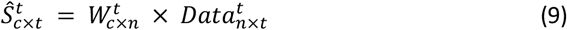

The spatial maps can be estimated by calculating the pseudo-inverse of the separating matrix *W*^*t*^.

## Methods

### Simulations

To guide the formation of our hypothesis regarding the effectiveness of sICA and tICA in identifying noise ICs, simulations were first performed, in which the fMRI data is simulated as a mixture of known components (ground truth). The same methodology has been used previously to simulate task-based fMRI (Calhoun et al., 2001). Each of the simulated datasets consists of five components of interest with known spatial and temporal patterns, as shown in Figure 1. Signals of the five components of interest are taken from an in vivo fMRI dataset used in this work (acquisition details to follow). The simulated dataset is decomposed into 50 ICs using the spatial ICA algorithm implemented in MELODIC, and for computational simplicity, only the initial 500 time points of each component are used. The spatial map of the simulated data consists of 500 voxels in a 2-dimensional matrix with 10×50 voxels (which can be conveniently divided into 5 sub-regions, each with a dimensionality of 10×10). In addition, one component with random spatial and temporal patterns are added to represent noise. Therefore, the mixing matrix *M* has dimensions of 500×6 (500 time samples and 6 components, i.e. 5 temporal components of interest and 1 noise component), and the components matrix S is a 6×500 matrix (500 voxels and 6 spatial components, i.e. 5 spatial components of interest and 1 random “noise” spatial patterns). Thus, the final resultant dataset has a dimensionality of 500×500. These components were used to produce four datasets to represent four different scenarios (Figure 1):

1. No spatial correlation and low temporal correlation; emulates random noise such as from thermal sources.
2. High spatial correlation and low temporal correlation; emulates spatially correlated but temporally asynchronous processes, such as visual activity and respiration in the occipital cortex (Birn et al., 2006; Golestani et al., 2015).
3. No spatial correlation and high temporal correlation; emulates spatially uncorrelated but temporally synchronous events, such as observed in different nodes of a brain network.
4. High spatial correlation and high temporal correlation: this is the most interesting and challenging case, in which we presume neuronal and vascular signal sources coincide both temporally and spatially (Bright et al., 2020).

**Figure 1.**
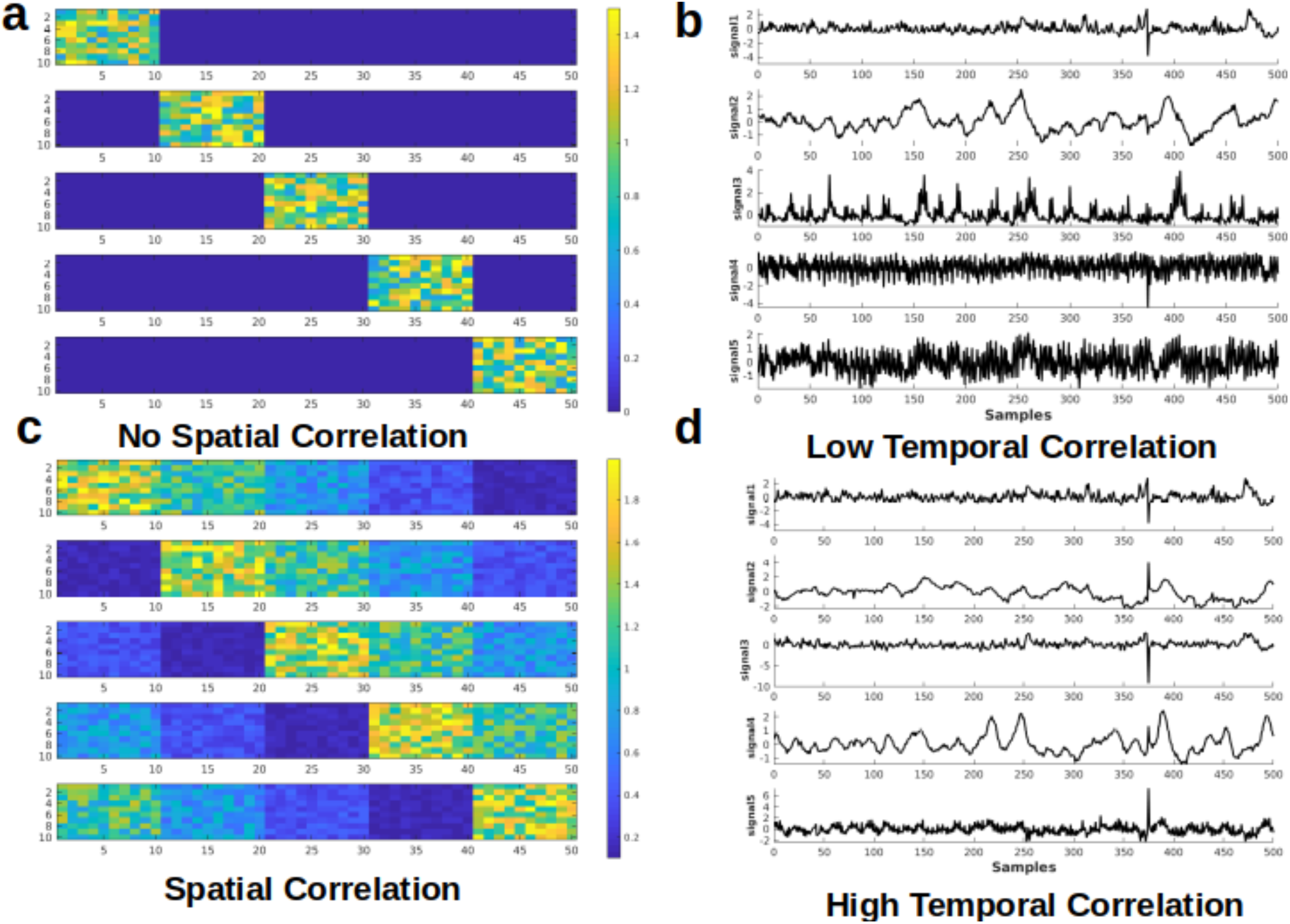
Simulated test cases. Examples of spatial (a & c) and temporal (b & d) components used for the simulation. Examples of the spatial maps of the five components are presented on the left. Each component has dimensions of 10×50. Examples of temporal signals of the five components are presented on the right. Each component has 500 time samples (time points). Panels a & b show the cases that the components have low correlation, whereas the panels c & d represent the cases where the components are highly correlated.

Regarding the component signals, for Scenarios 1 and 2, five signals are randomly selected from a set of 10 signals with the lowest mutual temporal correlation (the average absolute correlation among the 10 components = 0.0325), which reflects high independence among the temporal components. To model high temporal dependence for scenarios 3 and 4, five signals are randomly selected from a pool of ten highly mutually correlated signals (average absolute correlation among the 10 signals = 0.2701). Regarding the spatial components, for scenarios 1 and 3 (low spatial correlation), each signal is added to only one of the five sub-regions, whereas for scenarios 2 and 4 (high spatial correlation), each signal is added to all five sub-regions with different weightings. Examples of the components’ time-courses with low and high temporal correlation are shown in Figure 1b & 1d. To the best of our knowledge, this is a novel framework for determining the effectiveness of ICA-based methods for separating rs-fMRI-relevant signals contributions.

Datasets are generated by multiplying the spatial (*M*) and temporal (*S*) matrices. 100 datasets are generated for each scenario by varying the random noise, voxel values of the spatial patterns, and randomly selecting 5 out of 10 time-courses. Each dataset is then decomposed into its components using both sICA and tICA. The performance of the ICA algorithms is measured by comparing the spatial and temporal correlation between the 5 original and the 5 ICA-identified components. We realize that the assumption of temporally and spatially randomness for the noise component is an over-simplification in fMRI, but the goal of these simulations is to demonstrate the differential effects of tICA and sICA on temporally or spatially correlated signal components. The hypothesis is that while tICA should be better at separating temporally dissociated but spatially overlapping signal components, the converse should be true for sICA.

### Data acquisitions

19 healthy subjects (age = 26.5 ± 6.5 years) were scanned using a Siemens TIM Trio 3T MRI scanner with a 32-channel head coil. rs-fMRI scans were collected using simultaneous multi-slice GE-EPI BOLD (TR/TE = 380/30 ms, flip angle = 40°, 20 5-mm slices, 64×64 matrix, 4×4×5 mm voxels, multiband factor = 3, 1,950 volumes). During each scan, end-tidal CO_2_ pressure (PETCO_2_) fluctuations were passively monitored using a RespirAct™ system (Thornhill Research, Toronto, Canada). In addition, cardiac pulsation was recorded using the Siemens scanner pulse oximeter (sampling rate = 50 Hz), whereas the respiratory signal was recorded using a pressure-sensitive belt connected to the Biopac_TM_ (Biopac Systems Inc. California) at a sampling rate of 200 Hz. A T1-weighted anatomical image was also collected (MPRAGE, TR = 2400ms, TE = 2.43 ms, FOV = 256 mm, voxel size = 1×1×1 mm^3^).

### Preprocessing and ICA

The rs-fMRI processing pipeline includes motion correction, spatial smoothing (Gaussian kernel with 5mm FWHM), and high-pass filtering (> 0.01 Hz). We chose to estimate 30 ICs, as this is a typical number of components used in the literature that provides a good trade-off between providing a good representative of the fMRI data structure and making the analysis and interpretation more manageable (Vergun et al., 2016; Y. Wang & Li, 2015). For spatial ICA (sICA), fast ICA (Hyvärinen, 1999) is used to divide fMRI data into 30 spatial components. For temporal ICA (tICA), as is typical, the data dimension is first reduced to 100 components using sICA, and then tICA (Glasser et al., 2018; Smith et al., 2012) is performed on the 100 time-series to generate 30 temporal components. To assess the generalizability of our findings, we also obtained results when the signal was decomposed into 50 ICs (Supplementary Materials).

### Markers of noise: physiological variations and motion

We address the signal contribution by different noise types, categorized as:

- <u>Physiological fluctuations</u>, including PETCO_2_, RVT, and HRV, which have network structure and are spatially selective, but have temporal signatures that are distinct from those of neuronally driven BOLD signals. Heart-rate variation (HRV) is calculated as the average heart rate over a 4 second window (Chang et al., 2009). Respiratory-volume variability (RVT) is defined as the ratio of breathing depth to breathing period (Birn et al., 2006; Chang et al., 2009). PETCO_2_ is calculated by finding the peak PCO_2_ level in each breathing cycle and repeating over the entire tracing (Golestani et al., 2015). Subject-specific response functions for PETCO_2_, RVT, and HRV are obtained from the whole-brain global signal using the Gaussian-constrained maximum-likelihood deconvolution model (Falahpour et al., 2013; Golestani et al., 2015). The physiological signals are convolved with the corresponding response function.
- <u>Motion parameters</u>, including framewise displacement (FD), the spatial root-mean-square of the time series (DVARS), the slow variations (SVAR), and the six affine head motion parameters (3 rotations and 3 translations). Bulk-motion time series, whether fast or slow, are expected to exhibit statistical properties that differ from non-motion signal substrates both temporally and spatially. DVARS and SVAR are estimated using a MATLAB script (Afyouni & Nichols, 2018; Jenkinson et al., 2002). Specifically, DVARS is proportional to the sum of the squared framewise fMRI signal change and is weighted towards the fast portion of signal change. Conversely, SVAR is computed as the sum of the squared sum between consecutive fMRI frames and reflects the slow portion of signal change (Afyouni & Nichols, 2018; Jenkinson et al., 2002). The six affine motion parameters were generated using FSL’s MCFLIRT motion correction algorithm (Jenkinson et al., 2002). FD is calculated using FSL, as the sum of the absolute values of the derivatives of the six motion parameters.

### Evaluation methods

Evaluation of sICA and tICA for separating signal and noise are evaluated using the following evaluation approach, using the noise (physiological variability and motion) markers described earlier. In all cases, signal contributions associated with each noise marker are obtained by combining all ICs that are significantly correlated with each noise marker. Conversely, the remaining ICs are combined to synthesize the non-noise related contribution for each noise type respectively. Our methodology is detailed in Figure 2, and the evaluation rubrics are shown in Table 1.

1. **Noise identification: noise content in noise-correlated ICs** To assess the extent of a given IC indeed being mostly noise, the correlation between each noise time series and the time course of each of the 30 temporal/spatial ICs is calculated. To assess the significance of the correlations, a null distribution is generated by calculating the correlation between a specific component and 5000 permutations of the noise time course (to maintain the same power spectrum but with a shuffled phase). Noise-related ICs are defined as those that are significantly correlated with the noise (p < 0.05 Bonferroni-corrected for multiple comparisons). Ideally, the ICA should a) confine noise-related ICs to as few components as possible (low % of ICs identified as noise-related ICs), and b) produce noise ICs that are well correlated with the noise signals. Therefore, the performance of the ICA in noise-component identification is evaluated with the following parameters (Table 1):
  1.1. *<u>Percent of noise-related ICs:</u>* Percent of the components that have a significant correlation with the noise. A lower percent represents better performance.
  1.2. *<u>Noise-identification effectiveness ratio</u>*: Defined as the ratio of the average variance explained by noise in noise-related ICs (R^2^ between the noise time series and the time course of the ICA-identified noise-related components) divided by the average variance explained by noise in non-noise components (R^2^ between the noise time course and the time course of the non-noise ICs). Higher ratio represents better performance.
  **2. Noise removal:** Noise-related and noise-non-related ICs are combined separately to reconstruct “noise” and “denoised” datasets. To generate denoised datasets, all columns in the mixing matrix that are identified as noise components are replaced with zeros, following which the data is reconstructed by multiplying the mixing matrix to the components matrix. Similarly, “noise” datasets are generated by reconstructing data after replacing the non-noise columns of the mixing matrix with zeros. Ideally, successful noise correction should result in a “cleaned” dataset that contains high brain functional connectivity information. Moreover, the denoising approach should not remove excessive variance from the original data.

**Figure 2.**
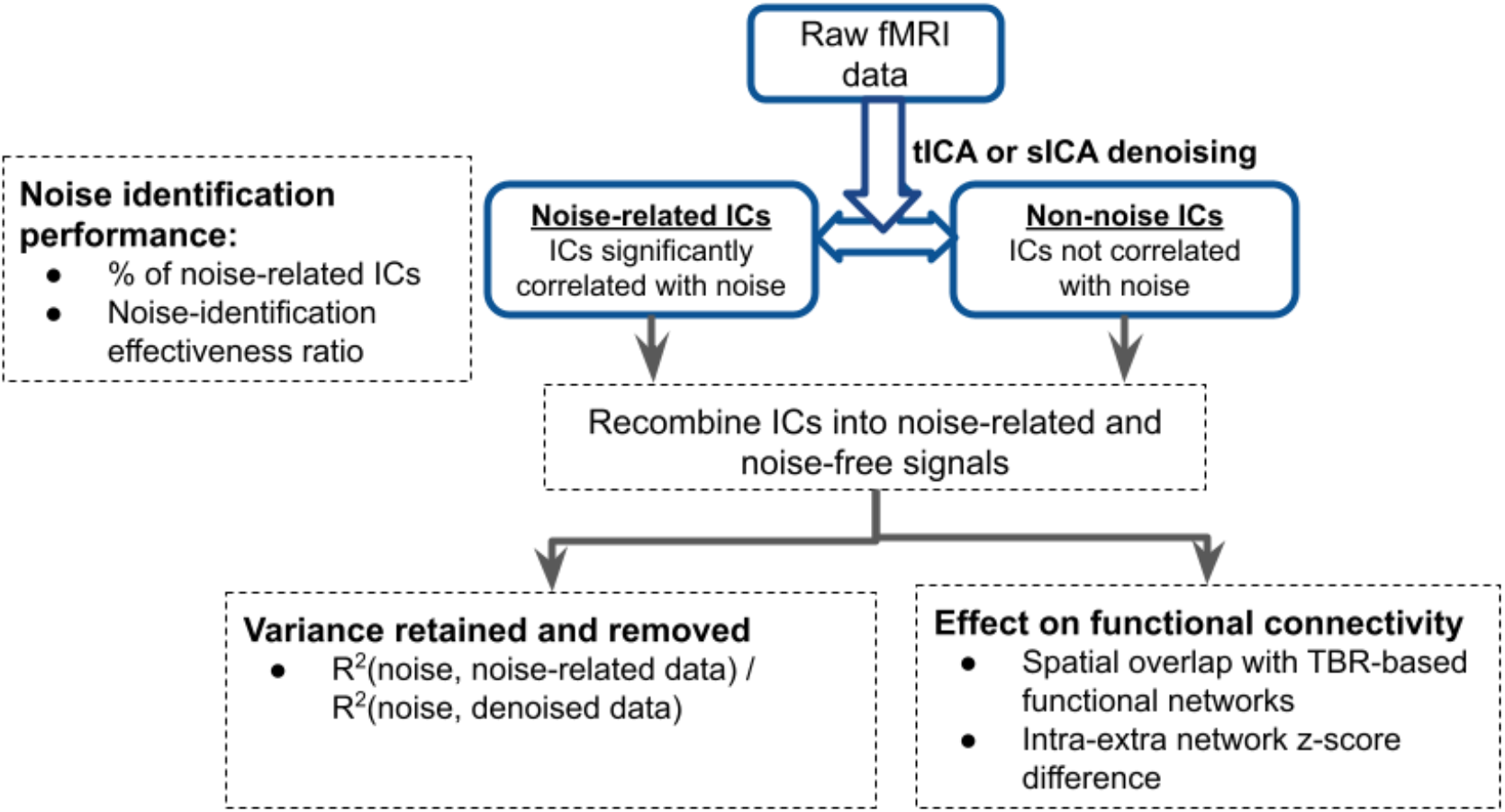
Overview of analysis pipeline for evaluating the performance of tICA and sICA. The performance of the two approaches are compared in three ways: (1) noise identification: the associated metrics are extracted from the results of the ICA step; (2) functional connectivity: these are extracted after combining noise-related and noise-free components to create noise and cleaned datasets ; (3) variance explained: the amount of variance (of the original data) that is removed from the pre-ICA signal (as noise) and how much is retained (noise-free), as well as the variance accounted for by the noise markers.

**Table 1.** Rubric demonstrating how to interpret the performance of noise identification (A) and correction (B) based on the presented data. A) A good noise identification consists of a few noise components with a strong correlation with the noise and a low correlation between the remaining components and the noise. On the contrary, a case of bad noise identification is when many components are correlated with the noise time series and the correlation between the noise and noise-related components is comparable with the correlation between the noise and noise-free components. B) Noise correction performance is evaluated by comparing the variance in the original data (before noise correction) and the variance in the cleaned and noise data explained by each noise signal. A good noise correction should reduce the R^2^ in the denoised data while having a high R^2^ in the noise data. Moreover, the Dice coefficient between rs-network templates and the components should stay the same or increase in the corrected data. Finally, **Δ**Z (the difference between the within-network and between-network Z-values) should not decrease after noise correction in the corrected data.

| Evaluation of Noise Detection Performance |  | Evaluation of Noise Removal Performance |  |
| --- | --- | --- | --- |
| % detected noise-related ICs | Ideally low | Denoising-effectiveness ratio:<br>Variance explained by noise markers in noise-related ICs /<br>Variance explained by noise markers in denoised signal | Ideally high |
| Noise-identification effectiveness ratio:<br>$R^2$(noise-related ICs, | Ideally high | Spatial overlap with known functional networks (Dice-non-noise) | Ideally high |
| noise markers) / $R^2$(non-noise ICs, noise markers) | | | |
| | | Intra-extra network connectivity difference ($\Delta Z$-non-noise) | Ideally high |

#### Variance retained and removed

In this step, we used the output from the ICA step to generate “noise” and “denoised” datasets with regard to different noise types (as described earlier). That is, for each noise time series, we identified ICs that are significantly correlated with it, and combined them to produce the noise-specific signal contribution for that noise type. Conversely, the remaining ICs are combined to create the signal contribution that is not related to that particular noise type (“denoised data” with respect to that noise type). To evaluate the effectiveness of noise removal, variance in the fMRI data explained by each noise source is compared before and after ICA-based noise removal. To this end, voxel-wise R^2^ between each noise source and the original fMRI signal was calculated and the R^2^ values were averaged across the brain. The same process was done for cleaned and noise datasets. The following measure is used in the evaluation:

- *<u>Denoising effectiveness ratio:</u>* defined as R^2^(noise signal, noise data) / R^2^ (noise signal, denoised data). Successful noise removal would lead to a decrease in the R^2^ between the noise signal and denoised data while R^2^ between the noise signal and noise data should be high. Therefore, we expect to have a high R^2^ ratio for a successful noise removal.

Furthermore, a composite noise dataset is created by a weighted summation of all the ICs correlated with any of the noise signals, and a noise-free dataset is created by a weighted summation of the remaining components. The weights for each component are based on the estimated mixing matrix. Voxel-wise R^2^ between the original fMRI dataset and the composite noise and noise-free datasets is calculated and then averaged across the brain. This shows how much of the variance in the original fMRI signal is removed by correcting for all noise sources.

#### Effect on functional connectivity

To assess the effect of ICA denoising on functional connectivity, template-based rotation (TBR) (Schultz et al., 2014) is implemented on “cleaned” datasets to generate resting-state connectivity (rs-connectivity) maps for each individual using Yeo’s seven resting-state network (rs-network) templates (Yeo et al., 2011), namely the visual, somatomotor, dorsal attention, ventral attention, frontoparietal and default mode networks. Specifically in TBR, functional volumes are described as a linear combination of network templates, and it is assumed that the network templates are meaningful segmentations of the rs-fMRI signal fluctuations. The first step of TBR is a spatial principal-component analysis of the fMRI data, resulting in mutually orthogonal principal components. These principal components are then mapped onto a network template using multi-regression. Thus, there is no requirement for the signals associated with individual network templates to be orthogonal. The same rs-fMRI image series could be used to map to multiple network templates, reflective of possible dependence amongst networks. The advantages for using TBR include that it provides more stable connectivity estimates as compared to traditional methods such as seed-based analyses, and that it offers a convenient means of incorporating the rs-fMRI network templates in our evaluation process.

Ideally, functional networks should be preserved in the “cleaned” images. As an example, group-average connectivity maps for the default mode network (DMN) are generated from TBR for the cleaned images. The following two measures are introduced to evaluate the presence of rs-networks in and “cleaned” data resulting from the ICA denoising stage:

2.1. *<u>Spatial overlap with known functional networks (Dice Coefficient)</u>*: Each network map generated using TBR is thresholded with a value that generates the maximum Dice coefficient with the functional-network template. The Dice coefficients are averaged across the six rs-networks (excluding limbic due to partial coverage and susceptibility noise). Ideally, concurrently high Dice coefficient from “cleaned” data demonstrates that the information about brain connectivity is preserved after noise correction (Table 1).
2.2. <u>Intra-extra network connectivity difference (</u>*<u>**Δ**Z</u>*): For each network, the “cleaned” data is mapped to individual network templates using TBR, as described earlier. Subsequently, the average z-values (connectivity score) taken from outside each network is subtracted from the average within-network connectivity. In a cleaned dataset we ideally expect to observe a greater difference between within-network and between-network connectivity.

### Statistical Test

Since the evaluation metrics are not always normally distributed, we used the paired-sample Wilcoxon signed-rank test to compare the metrics produced by the two methods (in addition to “no denoising”). To control for family-wise errors, only p-values of less than 0.01 are considered to be significant.

## Results

### Simulations

Results of the simulation are presented in Figure 3. For scenario 1 where the components have no correlation across the spatial or temporal patterns, sICA outperforms tICA in identifying spatial patterns, whereas tICA can better identify the temporal patterns of the components. In scenario 2, whereby the components are spatially but not temporally correlated, tICA displays better performance in identifying both spatial and temporal patterns of the components. In scenario 3, when the components are temporally correlated and spatially independent, sICA performs better in identifying the components’ spatial patterns. In Scenario 3, the performance of sICA and tICA in identifying the time-series of the components are comparable. In the scenario that the components are both spatially and temporally correlated, sICA can better identify components’ time series, while tICA can better identify components’ spatial patterns.

**Figure 3.**
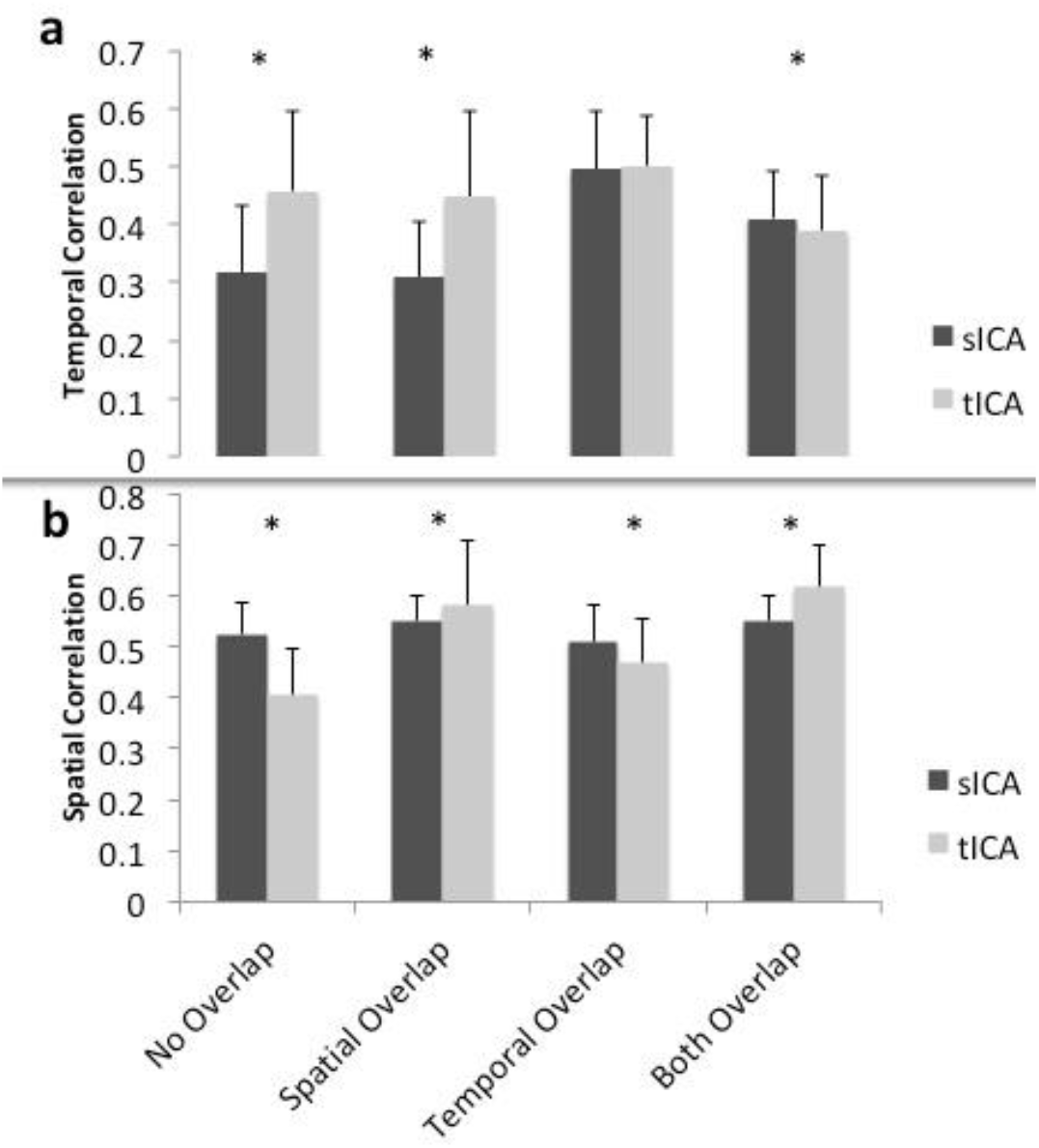
Simulated performance of sICA (black) and tICA (gray) in identifying signal components and spatial patterns. Performance is measured by calculating the correlation between the temporal (a) and spatial patterns (b) of the ground-truth and estimated components. Thus, a higher correlation indicates superior performance. When the components are spatially inter-correlated, tICA performs better, whereas when the components are temporally inter-correlated, sICA results are more favorable. Error bars show standard deviation. Significant differences are indicated by asterisks (p<0.05).

Overall, these simulations demonstrate that sICA performs better when the components are not spatially correlated and tICA performs better when the components signals have low temporal correlation, confirming or hypotheses.

### Experimental data

As described earlier, each raw data set is divided into 30 independent components (ICs) using either spatial (sICA) or temporal ICA (tICA). First, the performances of sICA and tICA in identifying noise components are compared using the two metrics explained in the first column of Table 1 (detailed in Methods). The performance of spatial and temporal ICA in noise removal is then compared using the three metrics in the second column of Table 1 (detailed in Methods). The distinction between the evaluations of noise identification and noise removal is that in the former case, we focus on the presence of noise contributions in the ICs of the original data identified as “noise-related”, whereas in the latter, we focus on the presence of noise contributions in the ICA-denoised data.

#### Noise identification

As shown in Figure 4 and Table S1, the Wilcoxson signed-rank test revealed significantly fewer tICA components than sICA components that are correlated with noise sources (indicated by blue asterisks). However, the noise-related ICs identified by tICA are more distinct from the non-noise related ICs, as indicated by a higher noise-identification effectiveness ratio (Fig. 4) for FD (red asterisk). In this respect (of the R^2^ ratio), sICA is only significantly advantageous in the cases of Y and Z translation (blue asterisks in the right columns).

**Figure 4.**
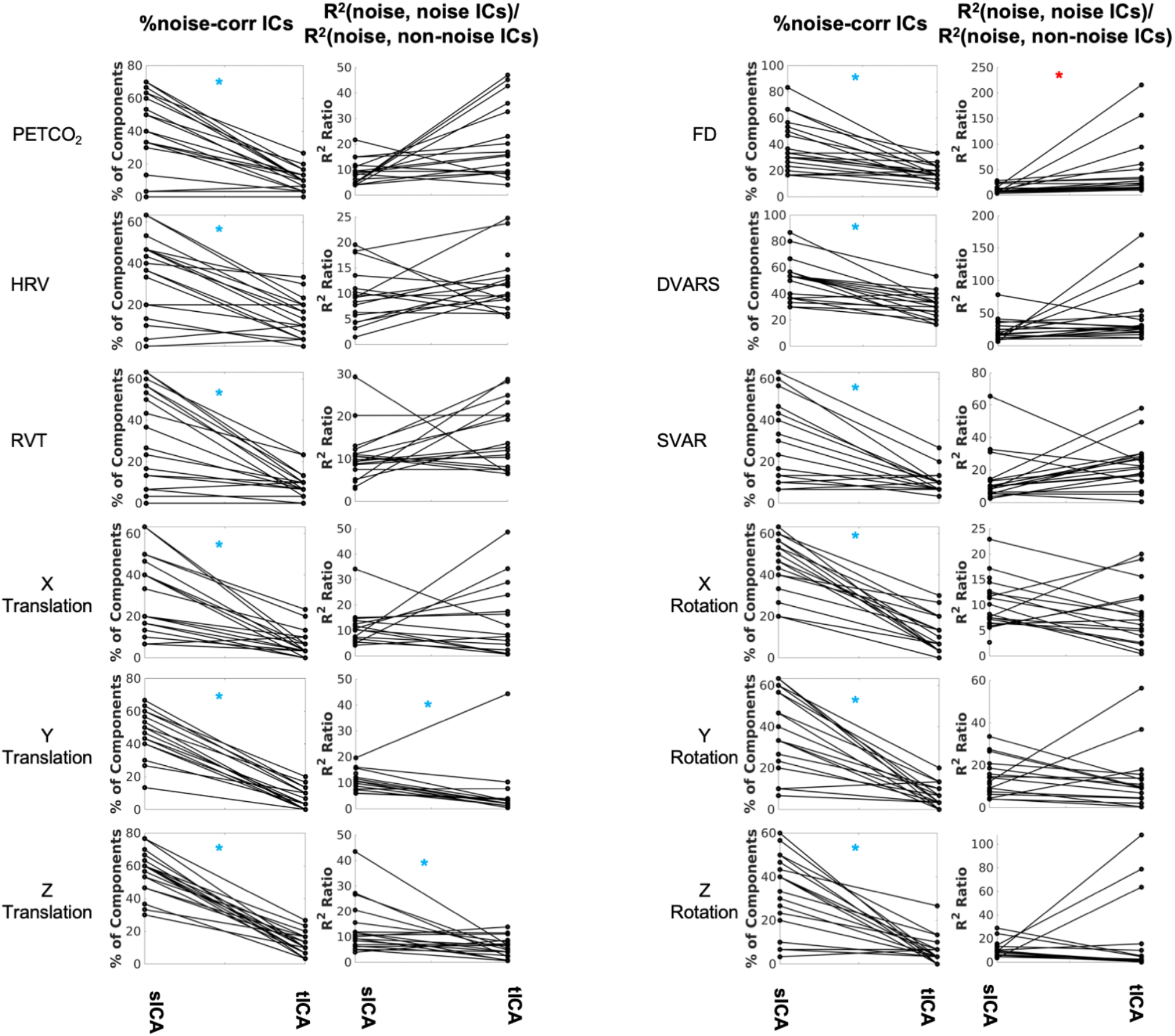
Comparing sICA and tICA in identifying noise components. Each line represents one subject. In each triplet of plots, column 1 depicts the percentage of components affected by noise signals, column 2 depicts the noise-identification effectiveness ratio, i.e., the ratio of R^2^ between noise signals and noise-correlated ICs over the average R^2^ between the noise signals and non-noise-correlated ICs. The significance of the changes is indicated in boldface in Table S1.

#### Noise removal

The noise components identified by both sICA and tICA have high shared variance (R^2^) with the noise sources, with a higher R^2^ ratio being indicative of higher denoising effectiveness (Table 1). By this metric, the performance of tICA for FD-related noise removal is superior to sICA (Figure 5, red asterisk), with the significance values summarized in Table S2. For the motion-realignment (translation and rotation) parameters, sICA performance is more consistent and significantly superior, specifically for Y and Z translation and rotation (blue asterisks, details found in Table S2). Similar findings pertain to the case of ICA producing 50 rather than 30 ICs (see Supplementary Materials).

**Figure 5.**
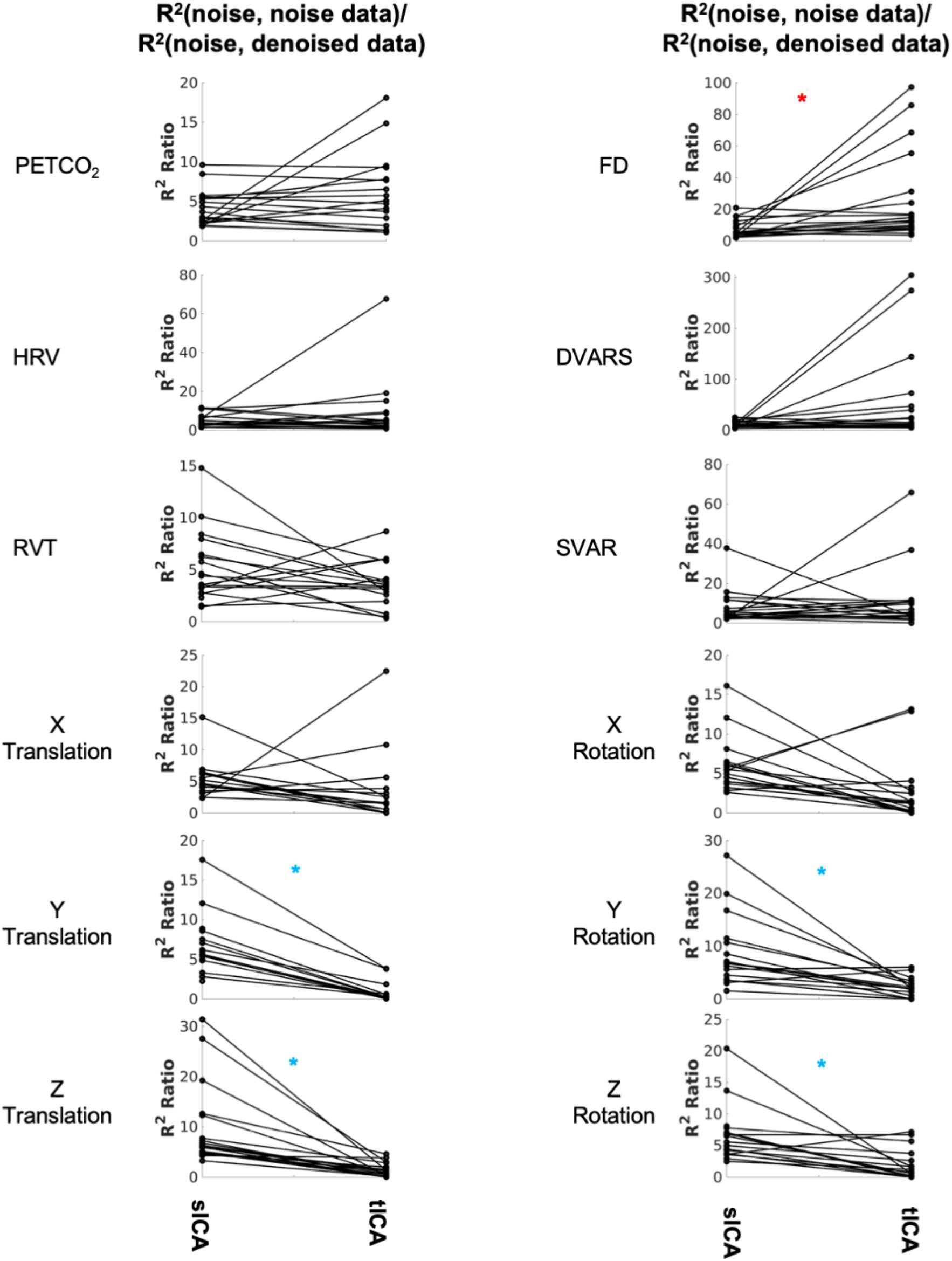
Comparing sICA and tICA in removing the effect of noise signals: the variance explained by noise signals. Each line represents one subject. In each column, each plot shows the ratio of variance accounted for by each noise/artifact source in the noise-related signal identified by sICA and tICA over the corresponding variance in the ICA-denoised signal. A higher R^2^ ratio indicates more successful denoising to some extent. Significantly higher values for sICA are indicated by the blue asterisk, whereas significantly higher values for tICA are indicated by the red asterisks. The significance of the changes is indicated in boldface in Table S2.

The effect of removing all noise sources is shown in Figure 6. Subjects are color-coded with different colors and symbols. In both cases, all ICs that are significantly associated with any noise source are considered noise-related and removed in the denoising step. Overall, tICA preserved considerably more variance of the original data. In some cases, sICA removed up to 80% of the variance in the original data.

**Figure 6.**
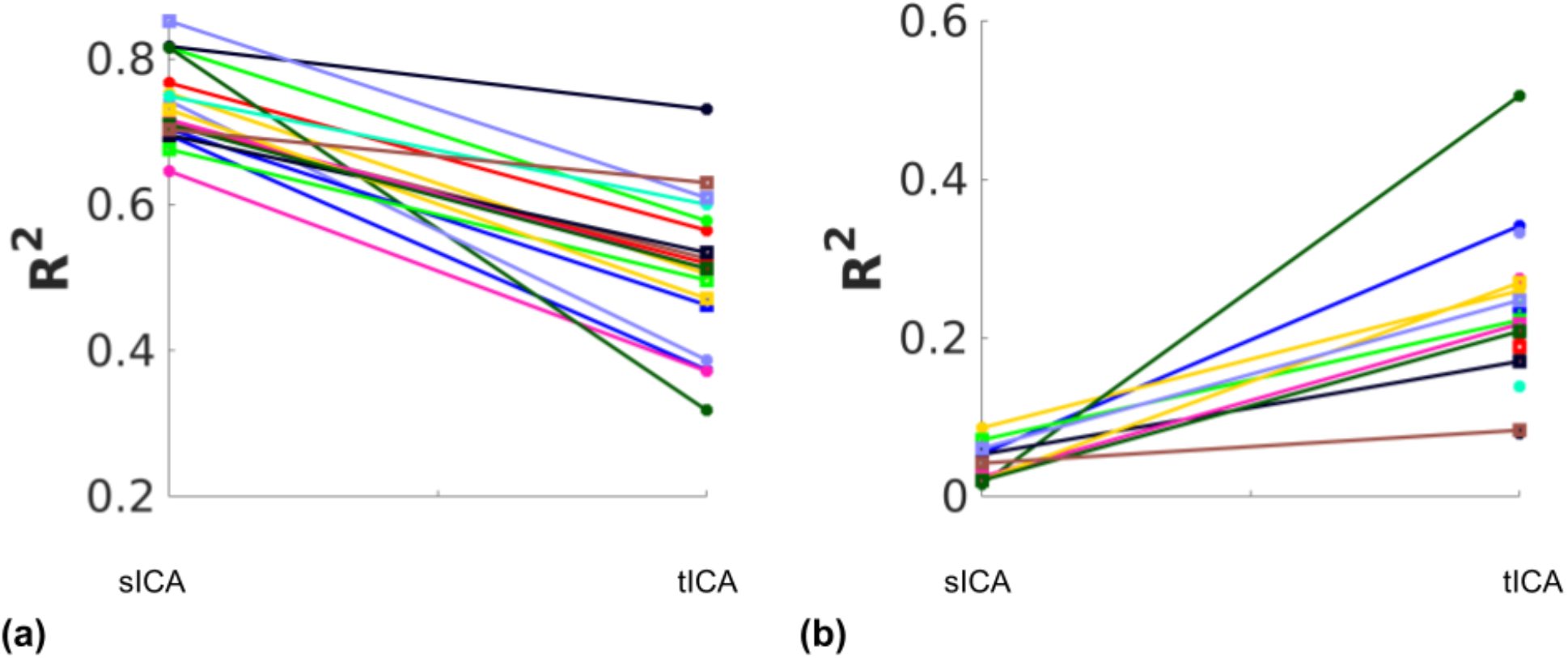
Total variance (R^2^) removed and retained by sICA and tICA. (a) The variance accounted for by the composite noise signal generated from the noise-correlated ICs. (b) The variance accounted for by the composite non-noise signal generated from the non-noise-correlated ICs. Each color represents one subject. The results show that sICA consistently removes more variance from the original signal than tICA.

To illustrate the influence on functional connectivity (FC), the DMN connectivity maps generated from corrected datasets are shown in Figures 7. The DMN generated from corrected data with tICA is more similar to the original DMN map, whereas the maps generated from corrected data with sICA have lower Z-values and, in some cases, missing nodes of the DMN network (e.g., dorsolateral-prefrontal node).

**Figure 7.**
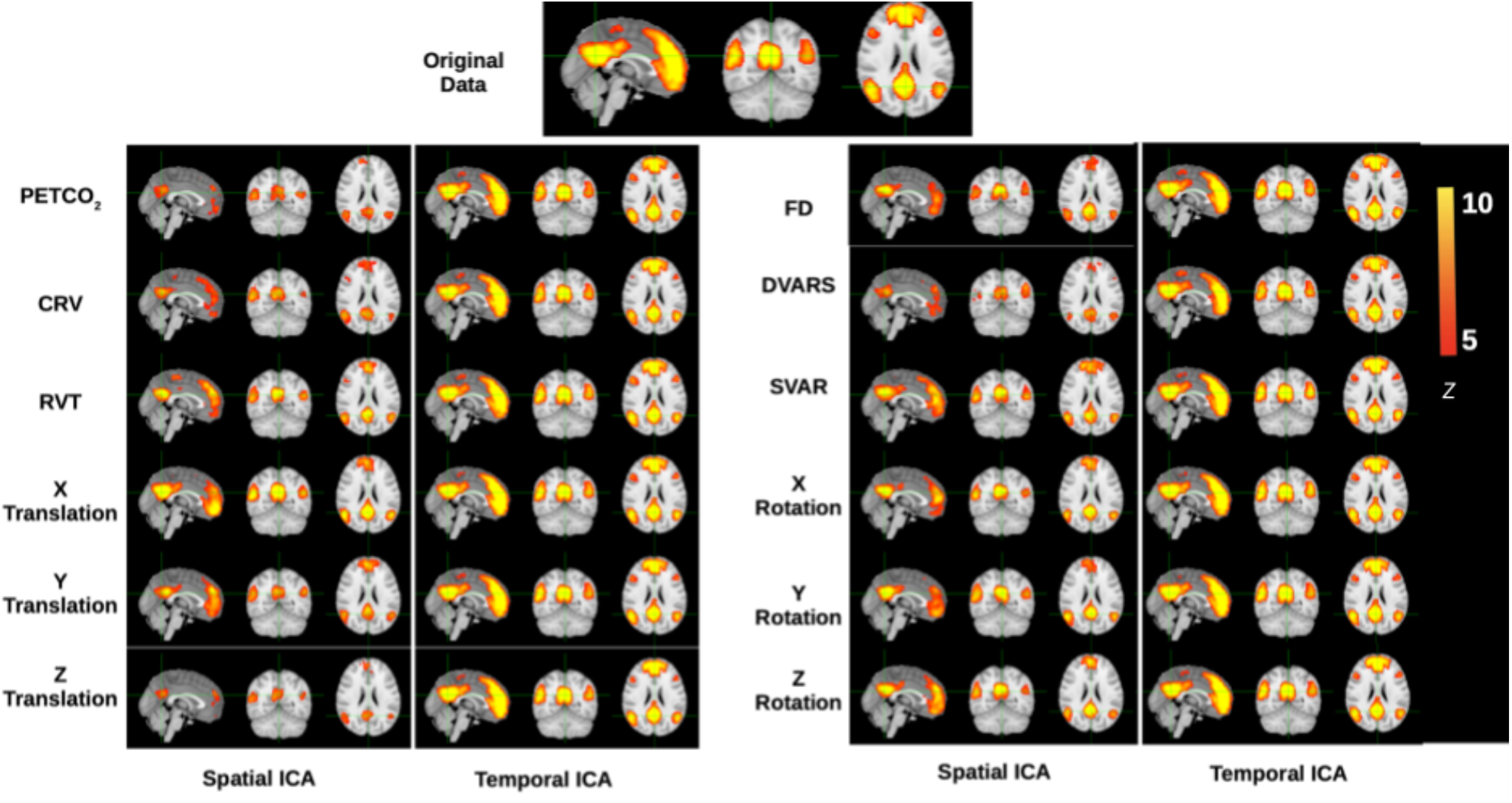
Functional-connectivity patterns associated with the denoised signals. Each row corresponds to the group-average TBR-based connectivity maps associated with signals that had specific noise correction applied to it. At the top is shown the DMN connectivity map obtained from the original data, and denoised data should maximally display DMN structure, which is generally stronger in tICA-cleaned signals.

To quantify the FC comparisons, the Dice coefficient was used to gauge the spatial similarity between each IC and template functional networks (Fig. 8 and Table S3). As mentioned previously, six networks were considered, namely the visual, motor, default-mode, dorsal attention, ventral attention and frontoparietal networks. In non-noise signals (those uncorrelated with each of the individual noise markers), tICA-denoised TBR results are shown to have significantly higher Dice coefficients, specifically after removing PETCO^2^, HRV, or RVT effects.

**Figure 8.**
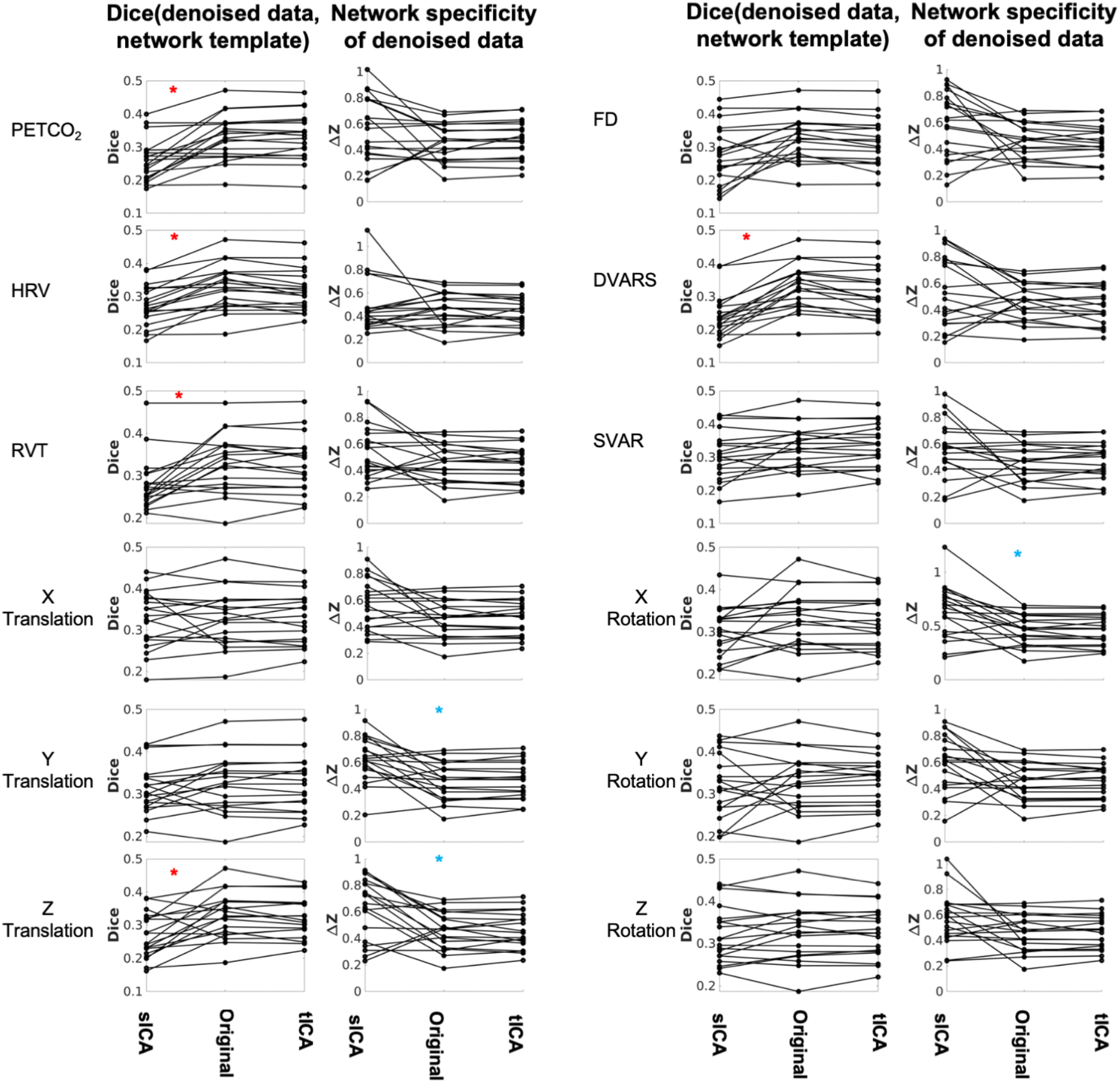
Comparing sICA and tICA in removing the effect of physiological signals: Spatial overlap of connectivity pattern with known functional networks (Column 1) and intra-extra network difference (Column 2). Each line represents one subject. Column 1 represents the average Dice coefficient between the rs-networks template and the resting-state network maps generated from denoised fMRI data. Column 2 represent functional specificity of each connectivity map as measured by calculating the average **Δ**Z between voxels within networks and those outside networks, and averaged over all networks. In each plot the ICA-denoising outcome is compared to the outcome from the original data (without ICA denoising). Significantly higher values for sICA are indicated by the blue asterisk, whereas significantly higher values for tICA are indicated by the red asterisks. The significance of the changes is indicated in boldface in Tables S3 and S4.

Network structure in non-noise related rs-fMRI signal contributions were assessed as the intra-extra network connectivity difference (**Δ**Z) (shown in Fig. 8 and Table S4) Compared to tICA, sICA denoising resulted in more variable changes in the network specificity. sICA exhibits greater inter-subject variability, as indicated by the spread of the Dice and network specificity metrics. The Dice coefficient (network-template overlap) is significantly higher for data after tICA-based removal of the effects of PETCO2, HRV, RVT, Z translation and DVARS (red asterisks), whereas network specificity (**Δ**Z) is significantly higher after sICA-denoising X rotation as well as Y and Z rotation (blue asterisks). Similar findings pertain to the case of ICA producing 50 rather than 30 ICs (see Supplementary Materials), demonstrating the generalizability of these findings.

## Discussion

ICA is a main-stream method of noise removal in resting-state fMRI (rs-fMRI) (Pruim et al., 2015; Salimi-Khorshidi et al., 2014). As ICA denoising can be purely data-driven, it circumvents the lack of physiological and motion recordings in many large-scale studies. However, to date, most ICA-related rs-fMRI studies have opted for spatial ICA (sICA) (Beckmann et al., 2005; Calhoun et al., 2001; Smitha et al., 2017), leaving temporal ICA (tICA) underexplored. We are cognizant of the rising use of accelerated rs-fMRI acquisitions (Demetriou et al., 2018; Lee et al., 2013; Preibisch et al., 2015), which is making tICA in rs-fMRI an increasing possibility. The effectiveness of tICA in identifying and removing the more global RVT effects in a group-wise tICA implementation has been shown (Glasser et al., 2018). In this study, we compare the performances of sICA and tICA for denoising rapidly sampled rs-fMRI data. Importantly, as we also have physiological and motion time series at our disposal, we are able to evaluate both types of ICA using these time series as reference rather than rely solely on more subjective evaluation. That is, noise-related ICs were identified based on significant correlation with noise parameters rather than based on spatial pattern or frequency distribution (Bhaganagarapu et al., 2013; Sochat et al., 2014). Furthermore, we use the available noise parameters to segregate the data into substrates driven by different noise types, namely physiological and motion.

In this study, although the spatial resolution (4×4×5 mm^3^) is lower than in studies such as the Human Connectome Project (Glasser et al., 2013), it was used such that we were able to achieve a high sampling rate (TR = 0.38 s), the intention being to help us avoid the brunt of aliasing high-frequency physiological noise in the f < 0.1 Hz band. Such spatial resolutions are not uncommon amongst legacy data that are still being actively analyzed (Gary H. Glover, 2011). It bears mentioning that we used a simulation-assisted approach to support our conclusions, an approach we have consistently embraced in our work (Chu et al., 2018; Yuen et al., 2019). In this study we assume that noise consists of all known signals in the frequency band < 0.1 Hz that have non-neural origins, including low-frequency physiological variability and head motion.

### Comparison of denoising methods: denoised resting-state fMRI signal content

tICA and sICA denoising behave very differently, although it is customary to precede tICA with sICA-based dimensionality reduction, as stated earlier (Glasser et al., 2018; Smith et al., 2012). The first main finding of this study is that tICA is likely to identify fewer noise ICs than does sICA for our fMRI data. Therefore, tICA results in a lower loss in degree of freedom and potentially preserves more neuronally meaningful signal contributions. At the same time, the noise-related ICs identified by sICA and tICA are similarly associated with the noise time courses, when normalized by the noise-series association with non-noise ICs. More specifically, as shown in Figure 5, while tICA performs significantly better for isolating the effect of FD, sICA performs better for isolating the influences of Y and Z affine motion realignment parameters. Furthermore, once all noise-associated ICs are removed, we found sICA removes much more variance from the original signal than tICA, as demonstrated in Figure 6; this is the second main finding of this study.

FD has a more global signature, as it is calculated from the whole-brain average signal, and acts like the summation of all affine motion parameters. On the other hand, the effect of the affine head motions can contain aspects that are localized at the edge of the brain with limited spatial overlap with brain networks and other signal sources. Thus, following the scenarios addressed in the simulations, tICA, as expected, performs better for isolating physiological noise (which share spatial distributions with brain networks), whereas sICA, as expected, performs better for noise types that have lower spatial overlap with brain networks. Nonetheless, we still need to assess whether these differences translate into equivalent performance differences in functional connectivity mapping. Lastly, the inter-subject variability of the performances of tICA and sICA are largely comparable, both exhibiting high levels of variability. This is also unsurprising, as the spatial signature of physiological noise varies greatly across subjects (Bianciardi et al., 2009; Chang & Glover, 2009; Falahpour et al., 2013).

### Comparison of denoising methods: resting-state fMRI connectivity

We found that sICA-denoised data is associated with lower network structure than tICA-denoised data, as shown in Figure 7 for the case of the DMN. When summarized across multiple brain networks, as the Dice coefficient reflects the degree of overlap between the network templates and the TBR maps of functional connectivity in the denoise data, a higher Dice coefficient in the denoise part of the signal is more ideal. Based on this metric, tICA denoising of physiological (PETCO_2_, HRV and RVT), DVARS and Z translation effects resulted in higher network integrity than sICA denoising (Fig. 8). Similar findings pertain to the case of ICA producing 50 rather than 30 ICs (see Supplementary Materials), demonstrating the generalizability of these findings. This is consistent with the finding that sICA denoising removes more variance from the data compared to tICA, part of which contributes to legitimate functional connectivity. These findings can be justified particularly as physiological (PETCO_2_, HRV and RVT) and DVARS share regions of influence (Bright et al., 2020; Y. Tong et al., 2015). Since sICA prefers spatial independence amongst ICs, it is less able to disentangle the effect of these noise signals from the underlying connectivity signals and therefore partially removes information about brain connectivity. On the other hand, sICA-denoised data produces higher network specificity in terms of X rotation and Y/Z translation, as these noise sources are likely to produce more spatially localized effects (Pruim et al., 2015).

Taken together, our findings suggest that there is real merit in considering a ICA-hybrid approach in which physiological and motion-related noise are each identified using tICA and sICA, respectively.

### Limitations

In this work we made a key assumption that noise in rs-fMRI should exhibit different statistical distributions from the signal. In reality, statistical distributions of physiological processes are not entirely distinct from those observed within functional networks. However, the inter-subject variability in these effects is very high (Chang & Glover, 2009; Golestani et al., 2015), creating large uncertainties as to the overlap with functional networks on a per-subject basis. The standard deviation of respiratory depth (RV) instead of RVT may be more robust against measurement artifacts (Glasser et al., 2018; Power, Lynch, Dubin, et al., 2019). As a result, the frequency occupancy of these signals can be leveraged to separate them from neuronally driven signals to some degree (Yuen et al., 2019). Nonetheless, this work presumes the currently dominant view that physiological processes are part of the “noise”. Further investigations are underway to verify that claim.

Furthermore, fMRI data typically has a much higher spatial than the temporal dimension. This leads to instability when applying tICA. To overcome this problem, it is inevitable to reduce the spatial dimensionality of the data. This is typically done using principal component analysis (PCA) (Boubela et al., 2013; Calhoun et al., 2001) or an initial spatial ICA (Glasser et al., 2018; Smith et al., 2012). In this study we performed an initial sICA-based data reduction, which is a common step in the tICA approach in fMRI (Glasser et al., 2018; Smith et al., 2012), to reduce the spatial dimension of the data to 100, which is a typical spatial space of the fMRI data (Craddock et al., 2012; Glasser, Coalson, et al., 2016). We did not test other dimension reduction methods or using sICA with a different dimension. Further studies are required to investigate whether different dimension reduction approaches would alter the findings.

Lastly, the spatial resolution of the current data is lower than used in the state-of-the-art studies (Glasser, Smith, et al., 2016; K. L. Miller et al., 2016). However, it is still representative of the numerous sets of legacy data (Kraus et al., 2020; Teipel et al., 2017), with an important advantage of critically sampling cardiac and respiratory noise peaks. This latter allows us a unique advantage to decipher the effects of low-frequency artifacts, although this results in the use of a lower than usual flip-angle (40 degrees). While such a low flip angle minimizes the contribution of physiological noise (Gonzalez-Castillo et al., 2011), given the loss in image SNR brought about by the short TR, the overall temporal SNR is no higher than at a higher flip angle (70-90 degrees). The TR-driven temporal-SNR can influence the performance of ICA and should be investigated as the next step.

## Acknowledgments

We acknowledge financial support from the Canadian Institutes of Health Research.

## Author Contribution Statement

Ali M. Golestani: conceptualization, data acquisition, data analysis, manuscript preparation.

J. Jean Chen: data acquisition, supervision of data analysis, manuscript preparation.

## Author Disclosure Statement

None of the authors have any competing financial interests to declare.

## Data Availability Statement

Data and code are available upon request from J. J. Chen.

## Funding Statement

This research was supported by the Canadian Institutes of Health Research (CIHR) Foundation Grant Program (FRN# 148398) and the CIHR Canada Research Chairs program.

## Notes

### Competing Interest Statement

The authors have declared no competing interest.

