## Supplementary Materials for "Performance of temporal and spatial ICA in identifying and removing low-frequency physiological and motion effects in resting-state fMRI"

**30 ICs**

The following tables quantify the results in Fig. 5-8 in the main body of the text.

**Table S1**. **Comparing sICA and tICA in identifying physiological signals, in terms of the percentage of ICs identified as noise** (assessed through correlations with various noise recordings), as well as the variance explained by noise in both noise and non-noise ICs. In most cases, sICA returned significantly more noise-related ICs.

|  | % of noise-correlated ICs **(mean (stdev))** | | | Noise identification effectiveness ratio **(mean (stdev))** | | |
| --- | --- | --- | --- | --- | --- | --- |
| **Type of Noise** | sICA | tICA | p | sICA | tICA | p |
| PETCO_2_ | 40.9 (23.1) | 10.4 (6.6) | **3.5e-4 (sICA > tICA)** | 8.5487 (5.0941) | 19.0421 (14.6422) | 0.0123 |
| HRV | 33.9 (19.1) | 13.9 (9.5) | **6.5e-4 (sICA > tICA)** | 8.7414 (5.5275) | 11.2408 (6.0054) | 0.1978 |
| RVT | 32.5 (22.5) | 9.1 (6.3) | **3.5e-4 (sICA > tICA)** | 9.8900 (6.4470) | 13.7392 (8.7571) | 0.0778 |
| FD | 38.8 (19.4) | 18.8 (7.2) | **3.8e-4 (sICA > tICA)** | 11.648 (7.882) | 44.193 (55.244) | **1.5e-4 (sICA < tICA)** |
| DVARS | 49.7 (15.9) | 30.7 (9.5) | **1.9e-4 (sICA > tICA)** | 22.473 (16.935) | 43.972 (41.707)) | 0.0442 |
| SVAR | 28.9 (19.0) | 10 (5.4) | **6.4e-4 (sICA > tICA)** | 14.043 (14.823) | 22.732 (13.924) | 0.0364 |
| X Trans | 31.4 (18.4) | 6.3 (6.6) | **1.5e-4 (sICA > tICA)** | 10.482 (6.651) | 11.234 (13.886) | 0.4445 |
| Y Trans | 45.1 (14.2) | 6.1 (6.3) | **1.3e-4 (sICA > tICA)** | 10.4707 (3.6640) | 4.4934 (10.0427) | **0.0025 (sICA > tICA)** |
| Z Trans | 56.3 (13.0) | 11.6 (7.0) | **1.3e-4 (sICA > tICA)** | 13.1483 (9.9661) | 5.7940 (3.7598) | **0.0062 (sICA > tICA)** |
| X Rot | 44.9 (14.8) | 10.7 (9.0) | **1.3e-4 (sICA > tICA)** | 10.1646 (4.8812) | 7.1438 (6.0727) | 0.0702 |
| Y Rot | 39.3 (19.3) | 6.5 (5.5) | **1.8e-4 (sICA > tICA)** | 13.4453 (8.5083) | 11.1438 (14.112) | 0.0766 |
| Z Rot | 33.9 (18.2) | 6.1 (6.3) | **2.5e-4 (sICA > tICA)** | 11.0832 (6.3606) | 15.6303 (31.298) | 0.1474 |

**Table S2**. **Comparison of sICA and tICA denoising outcomes in terms of the ratio of the variance explained by the noise signals in the noise and denoised datasets**. The former were assessed in ICs that were found to be significantly correlated to various noise sources, including physiological noise and head motion. A higher ratio is more ideal, as it indicates distinction between noise and non-noise ICs. While sICA and tICA denoising results are comparable for PETCO_2_, HRV, RVT, and SVAR, tICA-denoised results are associated with a lower R^2^ ratio in relation to the affine motion parameters (translation and rotation).

|  | **R^2^ Ratio (mean (stdev))** | | **p-values** |
| --- | --- | --- | --- |
|  | sICA | tICA | sICA vs Orig |
| PETCO_2_ | 3.9176 (2.3961) | 5.6215 (4.8223) | 0.5862 |
| HRV | 4.7284 (3.4429) | 8.3161 (15.2171) | 0.6292 |
| RVT | 4.9133 (3.5293) | 3.2077 (2.3493) | 0.0854 |
| FD | 7.4773 (5.3293) | 26.2195 (28.5924) | **0.0011 (sICA < tICA)** |
| DVARS | 11.0277 (5.8677) | 54.5557 (89.3401) | 0.0269 |
| SVAR | 7.6330 (8.3667) | 10.3812 (15.6485) | 0.8092 |
| X Trans | 5.0698 (2.7725) | 2.9982 (5.4151) | 0.0218 |
| Y Trans | 6.7259 (3.5113) | 0.6502 (1.1955) | **1.3e-4 (sICA > tICA)** |
| Z Trans | 9.5153 (8.0176) | 1.3199 (1.3525) | **1.3e-4 (sICA > tICA)** |
| X Rot | 5.8099 (3.3322) | 2.4107 (3.9237) | 0.0126 |
| Y Rot | 8.2444 (6.5795) | 1.9552 (1.8475) | **2.9e-4 (sICA > tICA)** |
| Z Rot | 6.5396 (4.2051) | 1.7212 (2.3699) | **5.4e-4 (sICA > tICA)** |

**Table S3**. Comparison of sICA and tICA denoising outcome, based on typical sICA results on the denoised signal. The evaluation is based on the average spatial overlap of ICs with known functional networks (quantified in terms of Dice coefficients) and the intra-extra network connectivity difference (𝚫Z). tICA-based results are associated with significantly higher Dice coefficients, specifically in signals that have been corrected for PETCO_2_, HRV or RVT. in these noise cases, sICA and tICA have comparable mean 𝚫Z values.

|  | **Dice coefficient (mean (stdev))** | | | **p-values** | | | 𝚫Z (mean (stdev)) | | | **p-values** | | |
| --- | --- | --- | --- | --- | --- | --- | --- | --- | --- | --- | --- | --- |
| **Type of Noise** | sICA | Original | tICA | sICA vs Orig | tICA vs Orig | sICA vs tICA | sICA | Original | tICA | sICA vs Orig | tICA vs Orig | sICA vs tICA |
| **PETCO_2_** | 0.259 (0.064) | 0.329 (0.069) | 0.332 (0.070) | **3.9e-4** | 0.4209 | **7.2e-4** | 0.554 (0.255) | 0.461 (0.143) | 0.479 (0.145) | 0.1024 | 0.0062 | 0.2598 |
| **HRV** | 0.273 (0.060) | 0.329 (0.069) | 0.317 (0.062) | **2.3e-4** | 0.0196 | **2.9e-4** | 0.487 (0.224) | 0.461 (0.143) | 0.450 (0.128) | 0.6791 | 0.0586 | 0.8405 |
| **RVT** | 0.279 (0.062) | 0.329 (0.069) | 0.326 (0.066) | **0.0050** | 0.4688 | **0.0038** | 0.540 (0.194) | 0.461 (0.143) | 0.452 (0.136) | 0.1446 | 0.2122 | 0.0702 |
| **FD** | 0.278 (0.088) | 0.329 (0.069) | 0.316 (0.070) | **0.0033** | **0.0019** | 0.036 | 0.582 (0.249) | 0.461 (0.143) | 0.447 (0.143) | 0.0442 | 0.1842 | 0.0196 |
| **DVARS** | 0.243 (0.065) | 0.329 (0.069) | 0.307 (0.072) | **1.3e-4** | **5.4e-4** | **2.5e-4** | 0.529 (0.265) | 0.461 (0.143) | 0.442 (0.157) | 0.1842 | 0.4688 | 0.990 |
| **SVAR** | 0.307 (0.075) | 0.329 (0.069) | 0.332 (0.066) | 0.0218 | 0.3341 | 0.0126 | 0.557 (0.208) | 0.461 (0.143) | 0.470 (0.139) | 0.0766 | 0.4688 | 0.1978 |
| **X Trans** | 0.327 (0.070) | 0.329 (0.069) | 0.330 (0.062) | 0.5461 | 0.4939 | 0.6292 | 0.559 (0.188) | 0.461 (0.143) | 0.465 (0.136) | 0.1075 | 0.5197 | 0.0836 |
| **Y Trans** | 0.312 (0.057) | 0.329 (0.069) | 0.331 (0.066) | 0.0990 | 0.9679 | 0.0298 | 0.623 (0.165) | 0.461 (0.143) | 0.470 (0.139) | **0.0033** | 0.1978 | **0.0048** |
| **Z Trans** | 0.267 (0.067) | 0.329 (0.069) | 0.323 (0.060) | **0.0029** | 0.3341 | **0.0043** | 0.626 (0.228) | 0.461 (0.143) | 0.461 (0.139) | **0.0079** | 1 | **0.0079** |
| **X Rot** | 0.302 (0.058) | 0.329 (0.069) | 0.324 (0.062) | 0.0702 | 0.3144 | 0.0766 | 0.631 (0.248) | 0.461 (0.143) | 0.455 (0.141) | **0.0043** | 0.3443 | **0.0033** |
| **Y Rot** | 0.319 (0.077) | 0.329 (0.069) | 0.330 (0.058) | 0.5732 | 0.6580 | 0.6874 | 0.592 (0.207) | 0.461 (0.143) | 0.462 (0.127) | 0.0176 | 0.6580 | 0.0126 |
| **Z Rot** | 0.321 (0.067) | 0.329 (0.069) | 0.329 (0.060) | 0.3144 | 1 | 0.3547 | 0.573 (0.200) | 0.461 (0.143) | 0.466 (0.132) | 0.1075 | 0.3981 | 0.0766 |

**50 vs. 30 ICs**

The following results correspond to the use of 50 instead of 30 ICs in the denoising and evaluation stages.

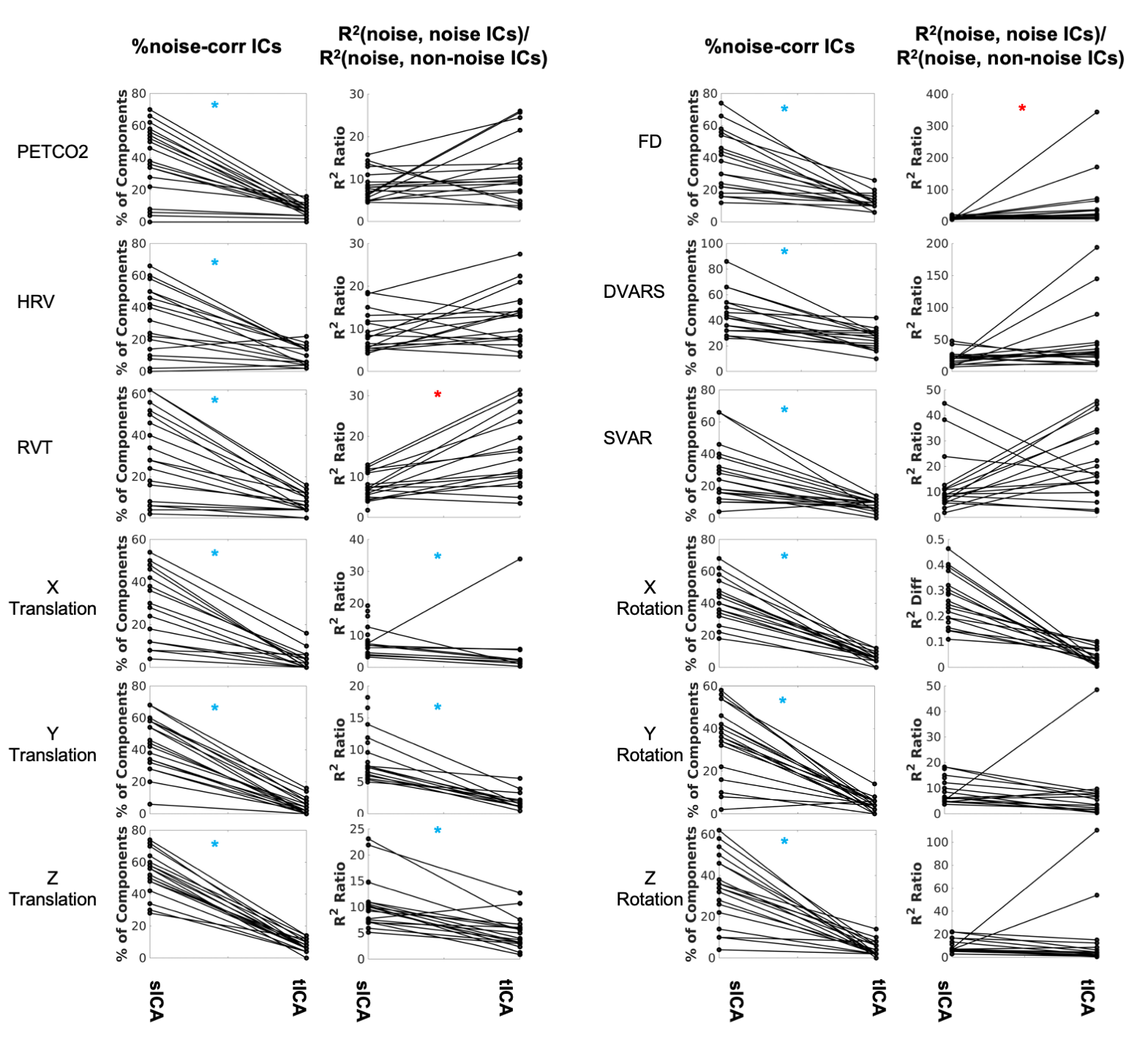

**Figure S1.** **Comparing sICA and tICA in identifying noise components.** Each line represents one subject. In each doublet of plots, column 1 depicts the percentage of components affected by noise signals, and column 2 depicts the ratio of the variance explained by noise in noise-correlated ICs over the variance explained by noise in noise-uncorrelated ICs. While tICA appears to have collapsed the noise contribution into fewer ICs, the variance explained by noise in these ICs is also higher for tICA than for sICA. Significance of the changes are indicated in boldface in Table S4.

**Table S4**. Comparing sICA and tICA in identifying physiological signals, in terms of the percentage of ICs identified as noise (assessed through correlations with various noise recordings), as well as the noise identification effectiveness ratio. In most cases, sICA returned significantly more noise-related ICs.

| Wilcoxon | % of noise-correlated ICs **(mean (stdev))** | | | Noise identification effectiveness ratio **(mean (stdev))** | | |
| --- | --- | --- | --- | --- | --- | --- |
| **Type of Noise** | sICA | tICA | p | sICA | tICA | p |
| PETCO_2_ | 38.1 (22.0) | 7.5 (4.0) | **1.9e-4 (sICA > tICA)** | 8.1304 (3.9654) | 11.3975 (7.8697) | 0.0707 |
| HRV | 32.8 (20.2) | 10.0 (5.8) | **4.9e-4 (sICA > tICA)** | 8.6705 (4.9220) | 12.6271 (6.3119) | 0.0364 |
| RVT | 30.3 (20.4) | 7.1 (4.6) | **1.9e-4 (sICA > tICA)** | 7.6696 (3.2716) | 14.4402 (9.8985) | **0.0033 (sICA < tICA)** |
| FD | 36.0 (18.9) | 12.5 (5.0) | **1.9e-4 (sICA > tICA)** | 10.8695 (4.3527) | 50.5246 (80.273) | **4.0e-4 (sICA < tICA)** |
| DVARS | 44.0 (15.7) | 25.2 (7.7) | **1.9e-4 (sICA > tICA)** | 21.0364 (10.391) | 42.9483 (48.746) | 0.1262 |
| SVAR | 27.1 (17.6) | 7.9 (3.7) | **3.2e-4 (sICA > tICA)** | 12.2793 (11.289) | 19.9556 (14.471) | 0.1165 |
| X Trans | 27.6 (17.6) | 3.4 (4.5) | **1.3e-4 (sICA > tICA)** | 8.6167 (4.5967) | 2.9591 (7.6856) | **0.0022 (sICA > tICA)** |
| Y Trans | 43.7 (16.9) | 4.4 (4.9) | **1.3e-4 (sICA > tICA)** | 8.7290 (3.8838) | 1.3964 (1.5049) | **1.3e-4 (sICA > tICA)** |
| Z Trans | 52.6 (13.0) | 7.5 (3.7) | **1.3e-4 (sICA > tICA)** | 10.7313 (4.8905) | 4.7700 (3.1841) | **3.4e-4 (sICA > tICA)** |
| X Rot | 43.1 (13.8) | 6.2 (3.2) | **1.3e-4 (sICA > tICA)** | 8.4524 (4.4321) | 5.4024 (4.9139) | 0.0486 |
| Y Rot | 34.7 (16.5) | 4.4 (3.3) | **1.7e-4 (sICA > tICA)** | 9.1185 (0.117) | 6.1251 (10.8024) | 0.0112 |
| Z Rot | 33.7 (16.7) | 4.4 (3.6) | **1.3e-4 (sICA > tICA)** | 9.6917 (5.7021) | 12.3637 (26.627) | 0.0218 |

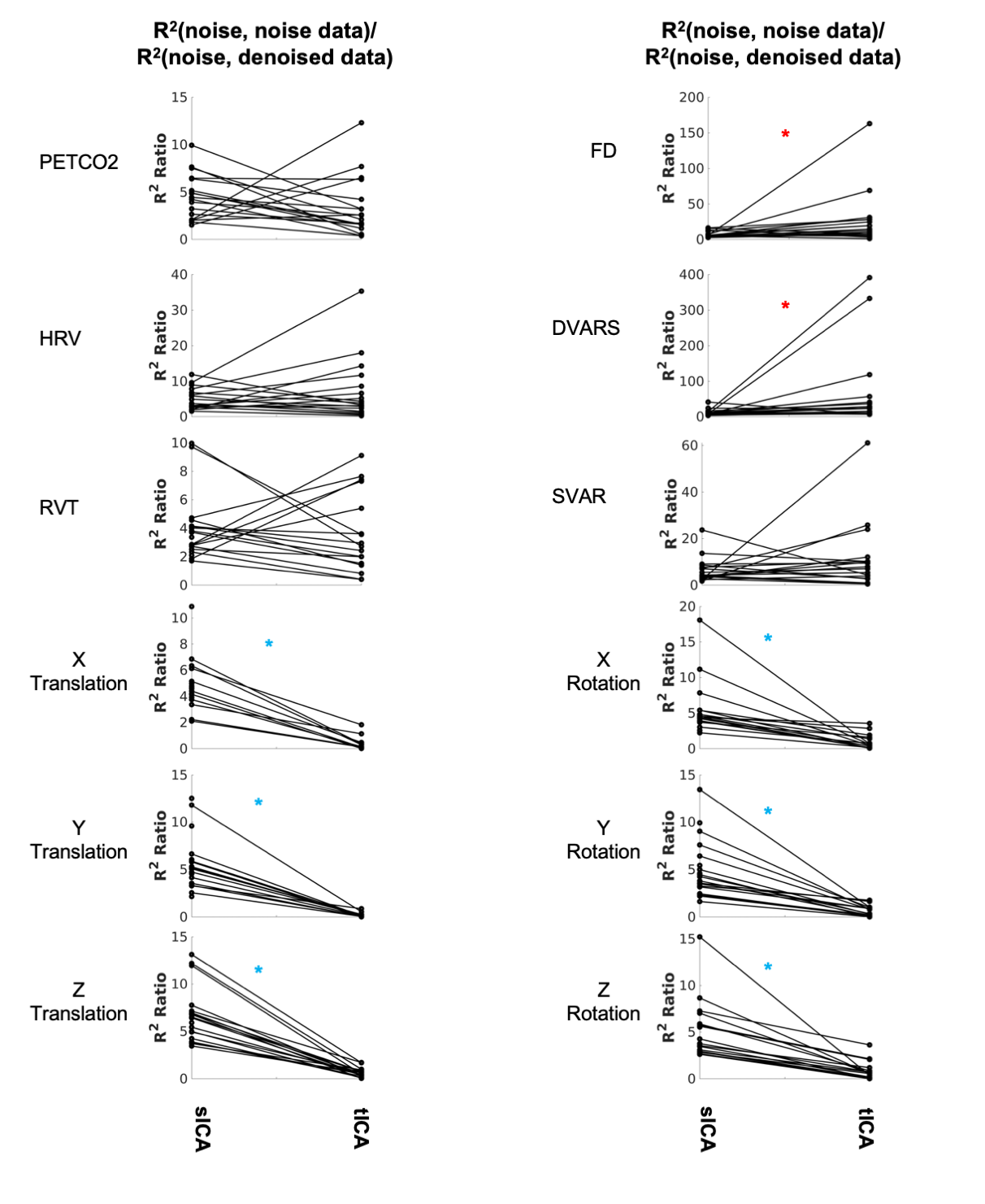

**Figure S2. Comparing sICA and tICA in removing the effect of noise signals:** the ratio of variance explained by noise signals. Each line represents one subject. tICA and sICA are associated with a similar level of residual noise for physiological noise and global motion parameters. The significance of the changes are indicated in boldface in Table S5.

**Table S5**. Comparison of sICA and tICA denoising outcome, in terms of the ratio of variance explained by noise in noise data over the variance explained by noise in non-noise data. A higher ratio is more ideal. As observed before, while sICA and tICA denoising results are comparable for RVT, FD, DVARS and SVAR, tICA-denoised results are associated with a lower R2 ratio in relation to the affine motion parameters (translation and rotation).

| Wilcoxon | **R^2^ Ratio (mean (stdev))** | | **p-values** |
| --- | --- | --- | --- |
|  | sICA | tICA | sICA vs Orig |
| PETCO_2_ | 4.2695 (2.5336) | 3.1508 (3.1146) | 0.1024 |
| HRV | 4.6915 (3.4429) | 6.6718 (15.2171) | 0.4939 |
| RVT | 3.8905 (2.2820) | 3.1981 (2.8466) | 0.3547 |
| FD | 7.1339 (3.9752) | 25.1539 (36.7872) | **0.0038 (sICA < tICA)** |
| DVARS | 11.8877 (8.5467) | 62.1699 (109.2605) | **0.0055 (sICA < tICA)** |
| SVAR | 6.2169 (5.1911) | 10.4773 (14.1911) | 0.6009 |
| X Trans | 4.8149 (1.8975) | 0.2436 (0.4727) | **1.3e-4 (sICA > tICA)** |
| Y Trans | 5.6245 (2.8367) | 0.1459 (0.2226)) | **1.3e-4 (sICA > tICA)** |
| Z Trans | 6.6367 (2.8914) | 0.5946 (0.4864) | **1.3e-4 (sICA > tICA)** |
| X Rot | 5.3878 (3.6466) | 0.7908 (1.0201) | **1.3e-4 (sICA > tICA)** |
| Y Rot | 5.0542 (3.0436) | 0.5273 (5650) | **1.3e-4 (sICA > tICA)** |
| Z Rot | 5.0483 (3.0550) | 0.7311 (0.9470) | **1.3e-4 (sICA > tICA)** |

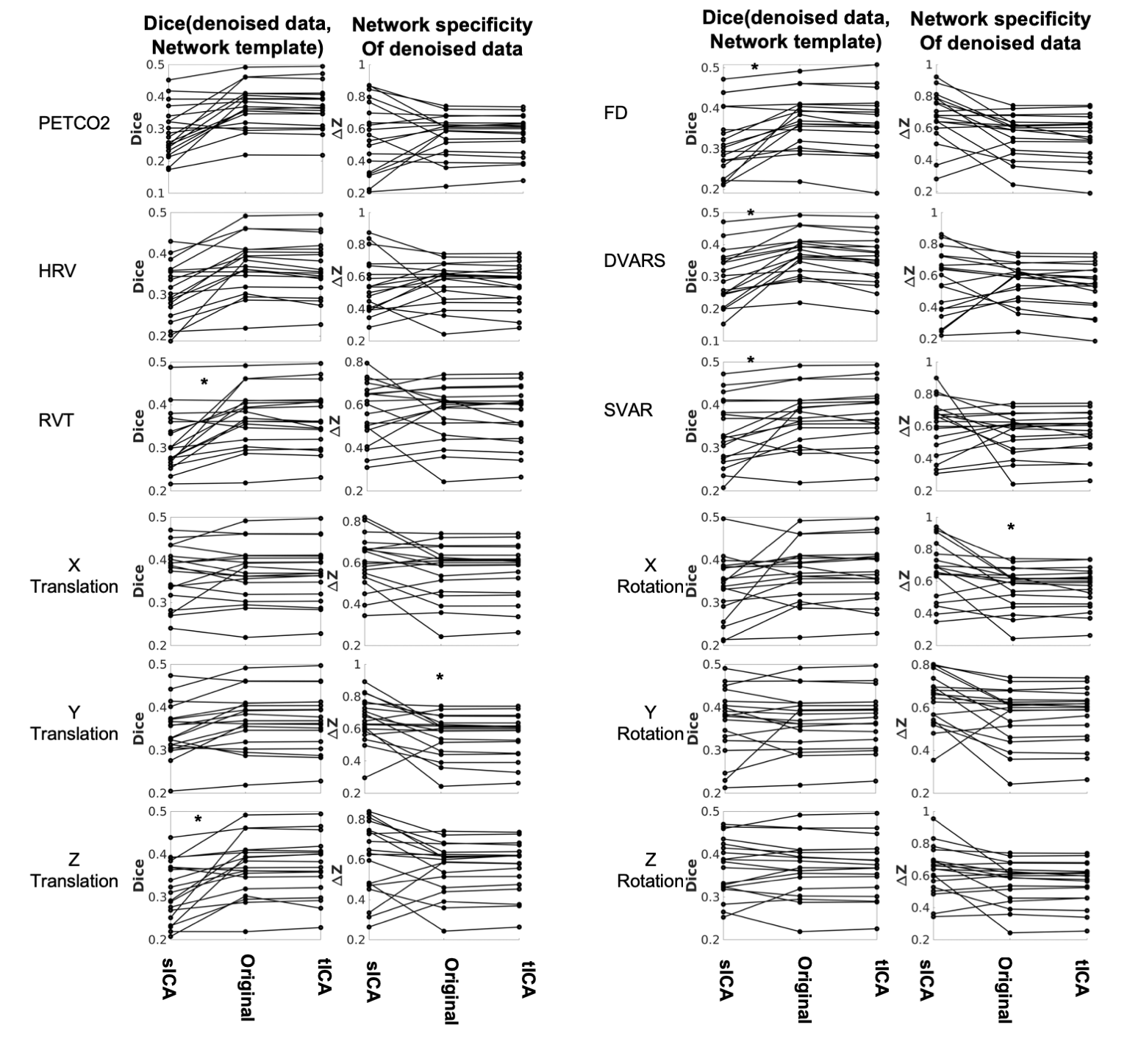

**Figure S3.** **Comparing sICA and tICA in removing the effect of physiological signals: Spatial overlap of connectivity pattern with known functional networks measured with Dice coefficient (Column 1), and intra-extra network difference (column 2).** Each line represents one subject. Each couplet of plots represent the average Dice coefficient between the rs-networks template and the resting-state network maps generated from corrected fMRI data (Column 1) as well as the average 𝚫Z computed between voxels within networks and those outside networks (Column 2). The significance of the changes are indicated in boldface in Table S6.

**Table S6**. Comparison of sICA and tICA denoising outcome, based on typical sICA results on the denoised signal. The evaluation is based on the average spatial overlap of ICs with known functional networks (quantified in terms of Dice coefficients) and the intra-extra network connectivity difference (𝚫Z). As observed before, in non-noise signals, tICA-based results display significantly higher Dice coefficients across nearly all rows. In most cases sICA and tICA have comparable mean 𝚫Z values.

|  | **Dice coefficient (mean (stdev))** | | | **p-values** | | | 𝚫Z (mean (stdev)) | | | **p-values** | | |
| --- | --- | --- | --- | --- | --- | --- | --- | --- | --- | --- | --- | --- |
| **Type of Noise** | sICA | Original | tICA | sICA vs Orig | tCA vs Orig | sCA vs tICA | sICA | Original | tICA | sICA vs Orig | tCA vs Orig | sCA vs tICA |
| **PETCO_2_** | 0.289 (0.079) | 0.370 (0.067) | 0.368 (0.068) | **6.3e-4** | 0.0990 | 9.7e-4 | 0.554 (0.218) | 0.561 (0.131) | 0.562 (0.125) | 0.8789 | 0.8092 | 0.9359 |
| **HRV** | 0.306 (0.069) | 0.370 (0.067) | 0.364 (0.068) | **4.6e-4** | 0.0242 | 8.4e-4 | 0.544 (0.168) | 0.561 (0.131) | 0.550 (0.130) | 0.3491 | 0.1842 | 0.5197 |
| **RVT** | 0.311 (0.067) | 0.370 (0.067) | 0.369 (0.068) | **5.4e-4** | 0.9359 | **0.0013** | 0.568 (0.139) | 0.561 (0.131) | 0.553 (0.133) | 0.6580 | 0.5461 | 1 |
| **FD** | 0.317 (0.081) | 0.370 (0.067) | 0.362 (0.074) | **5.4e-4** | **0.0025** | **0.0022** | 0.668 (0.161) | 0.561 (0.131) | 0.547 (0.144) | 0.0112 | 0.0641 | 0.0126 |
| **DVARS** | 0.294 (0.084) | 0.370 (0.067) | 0.352 (0.073) | **1.3e-4** | **3.4e-4** | **2.5e-4** | 0.553 (0.203) | 0.561 (0.131) | 0.536 (0.147) | 0.9039 | 0.0176 | 0.4688 |
| **SVAR** | 0.338 (0.075) | 0.370 (0.067) | 0.373 (0.070) | **0.0070** | 0.0218 | **0.0089** | 0.607 (0.165) | 0.561 (0.131) | 0.564 (0.128) | 0.7172 | 0.0641 | 0.5461 |
| **X Trans** | 0.365 (0.066) | 0.370 (0.067) | 0.371 (0.067) | 0.8721 | 0.1590 | 0.9679 | 0.606 (0.127) | 0.561 (0.131) | 0.562 (0.129) | 0.1262 | 0.3760 | 0.1075 |
| **Y Trans** | 0.345 (0.062) | 0.370 (0.067) | 0.370 (0.067) | **0.0089** | 0.6292 | 0.0141 | 0.656 (0.165) | 0.561 (0.131) | 0.559 (0.139) | **0.0062** | 0.9519 | **0.0062** |
| **Z Trans** | 0.313 (0.069) | 0.370 (0.067) | 0.371 (0.068) | **8.4e-4** | 0.0836 | **6.2e-4** | 0.604 (0.181) | 0.561 (0.131) | 0.565 (0.127) | 0.1590 | 0.1474 | 0.1590 |
| **X Rot** | 0.338 (0.072) | 0.370 (0.067) | 0.370 (0.069) | 0.0218 | 0.2432 | 0.0586 | 0.664 (0.173) | 0.561 (0.131) | 0.559 (0.126) | 0.0112 | 0.7782 | **0.0099** |
| **Y Rot** | 0.366 (0.077) | 0.370 (0.067) | 0.373 (0.065) | 0.5732 | **0.0062** | 0.8721 | 0.640 (0.123) | 0.561 (0.131) | 0.564 (0.127) | **0.0089** | 0.0766 | 0.0100 |
| **Z Rot** | 0.367 (0.067) | 0.370 (0.067) | 0.369 (0.066) | 0.8721 | 1 | 0.7782 | 0.632 (0.151) | 0.561 (0.131) | 0.558 (0.129) | 0.0364 | 0.3760 | 0.0196 |
